# Fishing with Two Lines: A Hybrid Approach to Spatial Transcriptomic Discovery

**DOI:** 10.64898/2026.01.07.698201

**Authors:** Arianna L. Williams-Katek, Saahithi Mallapragada, Evan D. Mee, Brandon K. Fischer, Laurie C. Eldredge, Gail H. Deutsch, Jonathan A. Kropski, Jennifer M. S. Sucre, Nicholas E. Banovich

## Abstract

Spatial transcriptomics faces a trade-off between the number of genes assayed and depth of per-gene sensitivity. We developed a ‘dual chemistry’ method that combines the high sensitivity of a 10X Genomics Xenium V1 custom panel (up to 480 genes) with the broad coverage of the Prime 5K panel (5001 genes) on a single tissue section. This involved co-hybridizing Prime and V1 probes and sequentially running the V1 and Prime decoding chemistries. Applied to a human lung tissue microarray, we observed high concordance between the V1 and Prime chemistries when run independently (on serial sections) and the dual chemistry runs. Overlapping genes (profiled on both V1 and Prime chemistries) showed similar expression patterns in the dual run demonstrating the fidelity of the assay. By combining information from both the V1 and Prime chemistries within the same cell, we retain more cells, gain valuable additional information, and enable both high sensitivity profiling and discovery.

## Introduction

The advent of commercially available platforms for profiling of gene expression changes at cellular or sub-cellular resolution, collectively known as spatial transcriptomics, is revolutionizing how researchers study tissue biology. Current commercially available instruments are largely split into two types of assays: imaging-based and sequencing-based. Imaging-based technologies generally depend on sequential cycles of in situ hybridization to decode transcript identity based on patterns of binding across cycles^1–3^. Sequencing-based technologies generally rely on a pull-down method of probe or transcript binding to a separate capture oligonucleotide^4^. The transcripts can then undergo library preparation and sequencing, then be mapped back to a barcoded coordinate grid based on the original tissue position on the slide. Across all platforms there is a tradeoff between breadth and depth. In general, lower plex imaging-based platforms provide high per gene sensitivity, while high plex imaging and sequencing-based platforms allow for broader coverage^5,6^. An idealized platform would deliver whole transcriptome coverage with high per gene sensitivity – similar to data from single cell genomics studies – but this has yet to be realized. The lack of such a platform leaves researchers to make difficult decisions about the value of breadth vs sensitivity in their studies, an issue that is further compounded by the lack of historical benchmarks due to the relative novelty of these approaches. Focusing on the 10X Genomics Xenium platform, we attempted to harness the sensitivity of the V1 chemistry (gene panels up to 480 genes in size) along with the breadth of the Prime chemistry (a predesigned panel of 5000 genes). Importantly, while the initial off-instrument sample preparation steps of the V1 and Prime chemistries are similar (see Methods) the on-instrument readout chemistry is substantially different. Thus, we present a method allowing for data generated from both the Prime human 5K panel as well as a V1 human 480 gene panel on the same slide. The V1 probes are spiked into the 5K probe mixture during sample prep. The probes are then decoded in successive Prime and V1 instrument runs. This results in a combined dataset that captures the breadth of the 5K panel alongside the sensitivity and depth of the V1 custom panel.

## Results

Three serial slides of a 17-sample (human lung) TMA were prepared using custom V1 probes, human Prime 5K probes, and a combination of the two termed the “dual” slide. The dual slide contained both sets of probes and was loaded on two unique Xenium runs, one V1 and one Prime, to create a dataset of cells with transcriptomic information from both panels (Fig. 1a). The V1 solo and Prime solo slides served as technical performance controls for the same 17 samples if only one chemistry was used. For the dual slide, the V1 chemistry was selected to load on the instrument first as it has a shorter runtime. First comparing the V1 chemistry on the solo vs dual preparation, we found both approaches to segment a similar number of cells (1,111,107 solo vs 1,107,543 dual) and decode a similar number of transcripts (221,439,804 solo vs 218,969,052 dual; Table 1). Importantly, the variance between the solo and dual V1 data was consistent with previously observed levels of slide-to-slide variation between serial sections (Table S1). Turning next to the Prime solo vs the Prime dual preparation (run after the completion of the V1 dual run), we again found a similar number of segmented cells (1,100,358 solo vs 1,115,068 dual) but a drop in the overall number of decoded transcripts (210,218,833 solo vs 187,967,974 dual; Table 1). This suggests running a V1 run before the Prime run modestly decreases the sensitivity of the Prime assay.

**Table 1:**
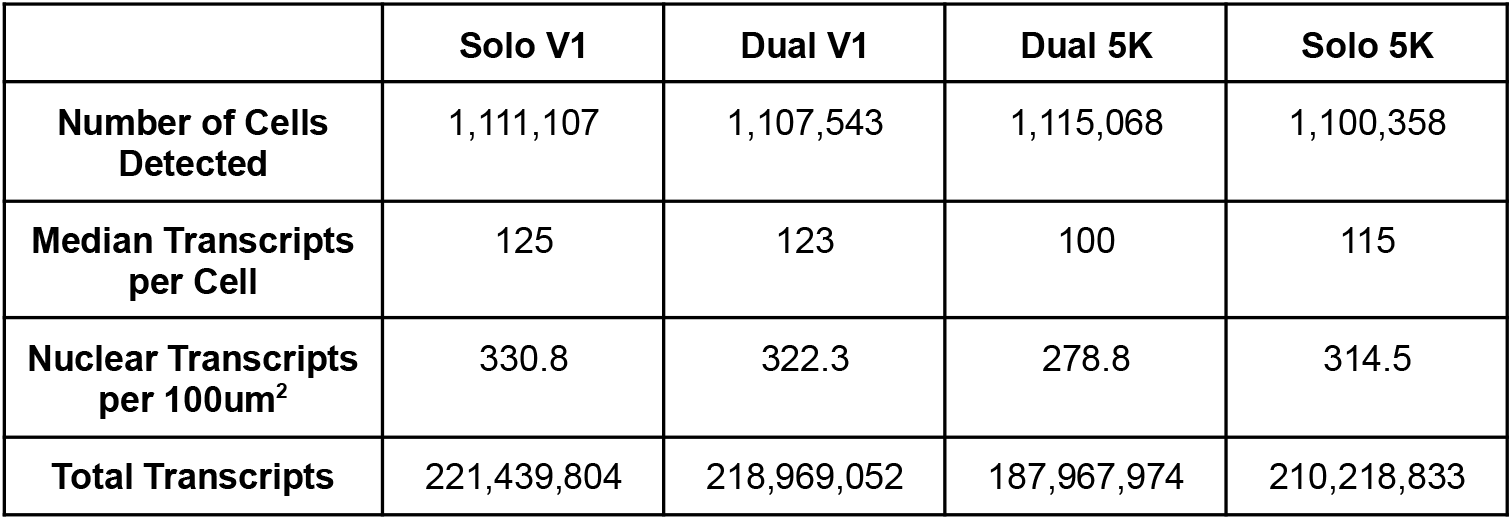
Slide-level transcript detection metrics. A table containing per slide segmented cell numbers, median transcript count per cell, average transcripts overlapping segmented nuclei per 100um^2^, and the total transcripts detected across the full slide.

**Figure 1:**
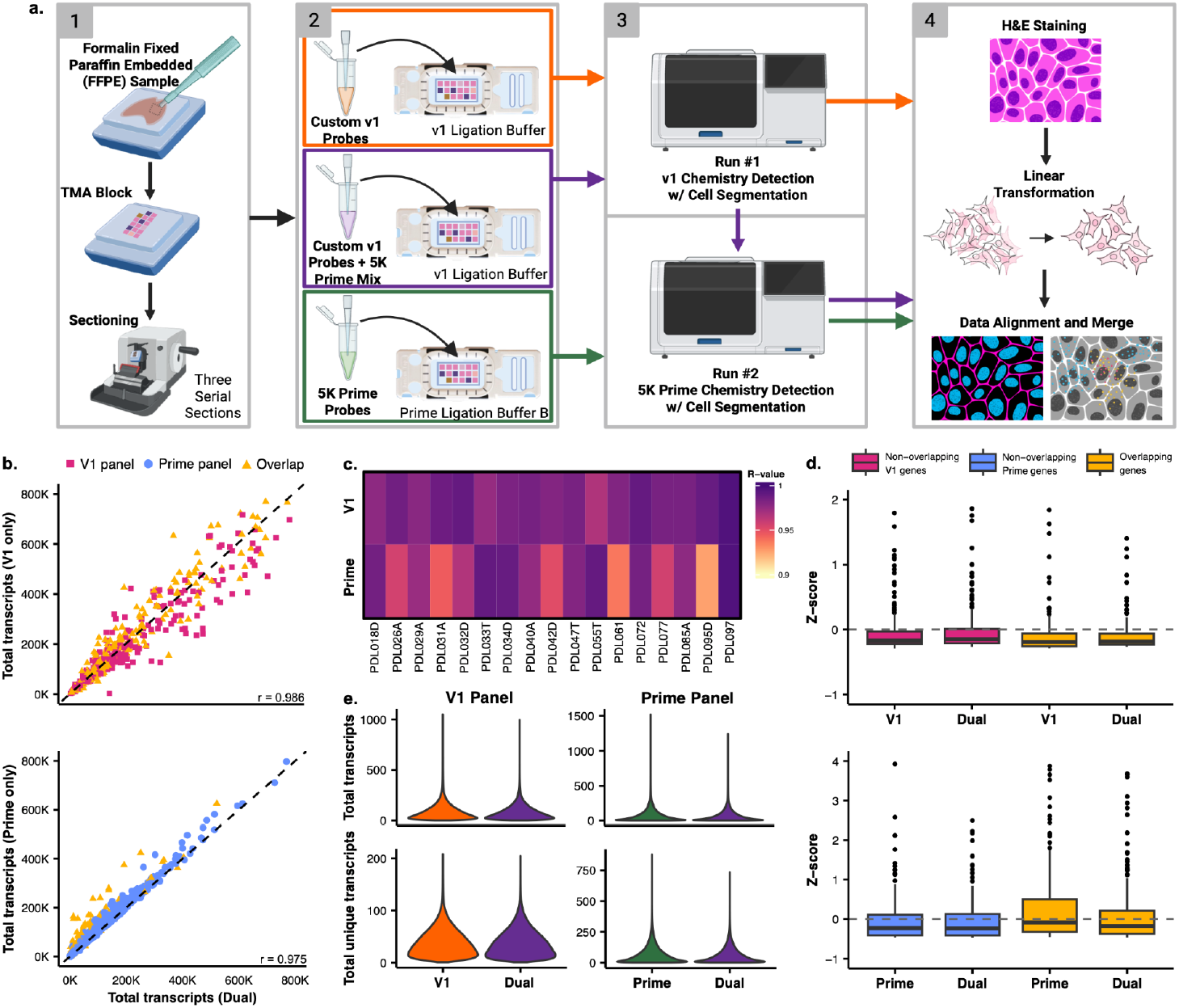
Evaluation of Transcript Quantification Consistency and Performance Across V1 and Prime Panel Assays. a) Schematic representation of the study design. b) Scatterplots that show the correlation of raw per-gene transcript counts across all genes detected in the V1 panel (top) and the Prime panel (bottom). The x-axis represents total gene expression from the dual assay (V1 + Prime or Prime + V1), while the y-axis represents total gene expression from the solo assay (V1 only or Prime only). Genes are colored by panel specificity: V1-specific genes (pink), prime-specific genes (blue), and genes overlapping both panels (yellow). c) Heatmap visualizing the R-value derived from correlating the transcript counts from the solo runs with the dual-run transcript counts for 17 infant lung samples. d) Boxplots that display the z-score distribution for three gene sets: the 239 genes overlapping between the V1 and Prime panels, the non-overlapping V1-specific genes, and a random subset of 239 non-overlapping Prime-specific genes. Genes are colored by their group: Overlapping (yellow), non-overlapping V1-specific (pink), and non-overlapping prime-specific (blue). e) Violin plots of the distribution of total transcripts and total unique transcripts for the three main run types: V1 only, prime only, and the combined dual run.

In this study, the V1 panel was designed specifically for use with human lung tissues whereas the Prime panel was designed to encompass gene expression in a variety of human tissues. The V1 panel detects a total of 480 genes and the Prime panel detects a total of 5001 genes. A 239 gene target overlap exists between the two panels allowing us to test for impacts of the same gene being targeted in both panels (Tabs. S2 - S3). Raw transcript count correlations were examined between the solo and dual slides of each chemistry. The solo V1 and dual V1 runs were highly correlated (r = 0.986; Fig. 1b) with no obvious impact on genes overlapping the V1 and Prime chemistries. The Prime solo and dual runs were also highly correlated (r = 0.975; Fig. 1b), however, in the genes targeted by both Prime and V1 we observed an exacerbated shift towards lower counts in the dual run compared to the solo run (Fig. 1b). While these correlations were performed at the slide level, each slide contained samples from 17 individuals. At the sample level, high correlation between solo and dual runs are maintained across both chemistries – with none dropping below r = 0.9 – although higher sample to sample variance is observed with the Prime chemistry (Fig. 1c, Figs. S1 & S2, Table S4).

We were concerned that genes targeted with both chemistries may inhibit one another. While none of the Prime and V1 probes were expected to bind the same target sequence, sensitivity could still be affected by off-target binding or steric effects of ligation and amplification machinery. To determine whether genes targeted by both V1 and Prime could be driving some differences in performance between the Prime solo and the Prime dual, we quantified sensitivity differences between overlapping and non-overlapping genes. Briefly, within each run (V1 solo, V1 dual, Prime solo, and Prime dual) we calculated Z scores of expression across the slide. We then examined the Z scores of the 239 overlapping genes compared to a random subset of 239 non-overlapping genes. If competitive binding drives a difference in sensitivity, we would expect to see a difference in the Z score distribution between the overlapping genes in the solo and dual runs but not the random subset of non-overlapping genes. Indeed, while we observed similar distributions for V1 solo and dual runs (Fig. 1d), we found the Z scores for the Prime solo to be higher than those for the Prime dual – for those genes that overlap between chemistries (Table S5). This is consistent with the shift towards lower counts seen in genes targeted by both chemistries observed in Fig. 1b, suggesting that competitive binding between V1 and prime probes is modestly reducing sensitivity in the Prime dual run but not the V1 dual run.

Another measure of run quality is the total number of transcripts detected on the slide. This metric remained high across all three slides, with detection only dipping under 200,000,000 transcripts on the Prime dual run. When compared within each chemistry, the dual slide shows no major variation in the distribution of total transcript per cell or total unique transcript per cell counts (Fig. 1e). This remains true on the per sample level, with multiple samples even outperforming at the cell level on the dual slide when compared to the solo slide in the V1 (Fig. S3) and the Prime chemistry (Fig. S4).

The key benefit of running both Xenium chemistries on the same slide is for co-detection of the higher sensitivity V1 and the broader reaching Prime panel in the exact same cells. While an adjacent section provides a decent proxy, it is still a 5um offset that then relies on the alignment of non-matched nuclei and is further complicated by the fact that cells exist in a 3D space with cytoplasmic overlap at that 5um scale. In order to combine data from the two runs of the dual slide, the transcripts first need to be put in a common cell coordinate system. In order to do this, a single segmentation mask must be applied to both runs. In this case we chose the mask from the V1 run. However, a linear transformation had to be applied to the original V1 run segmentation masks to align them with the slightly shifted cell placement from the Prime run (Fig. 2a). The shift occurred due to small technical variation that can occur when exchanging the dual slide from a V1 Xenium cassette and into a Prime Xenium cassette, as well as any slight changes to the exact placement of the cassette on the instrument stage during the second run leading to very small offsets. We expect this value to be variable and will thus require some amount of correction on any future slides run using this method. After the linear transformation, the detected transcripts falling within those bounds could be united under the same cell-id for both the V1 and Prime runs. Cells were then filtered using the following parameters to eliminate cells that would not meet normal thresholds for downstream analysis: n_counts >= 50, n_genes >= 5, cell_area >= 5 & <= 140, nucleus_area >= 3. These cutoffs were used to filter the 1,081,011 cells using transcripts only from the V1 panel, transcripts only from the Prime panel, and transcripts from the combined dataset. Of the four filter parameters, we found that n_counts had the greatest impact on cell retention. Using the combined transcripts left a greater number of retained cells, post-filtering, compared to the V1 or Prime panel transcripts alone (Fig. 2b). 75.5% of cells met the thresholds in the combined dataset, 265,161 cells lost and 815,850 retained. 59.6% of cells met the thresholds using V1 transcripts only, 436,942 cells lost and 644,069 retained. Less than half of the cells at 47.2% met the thresholds using Prime transcripts only, 570,806 cells lost and 510,205 retained. This trend of the dual data set retaining the most cells and the Prime panel alone retaining the least was consistent across all samples, with one exception in PDL034D where the Prime panel alone retained 149 more cells than the V1 panel alone with both still falling short of the combined panel (Table S6).

**Figure 2:**
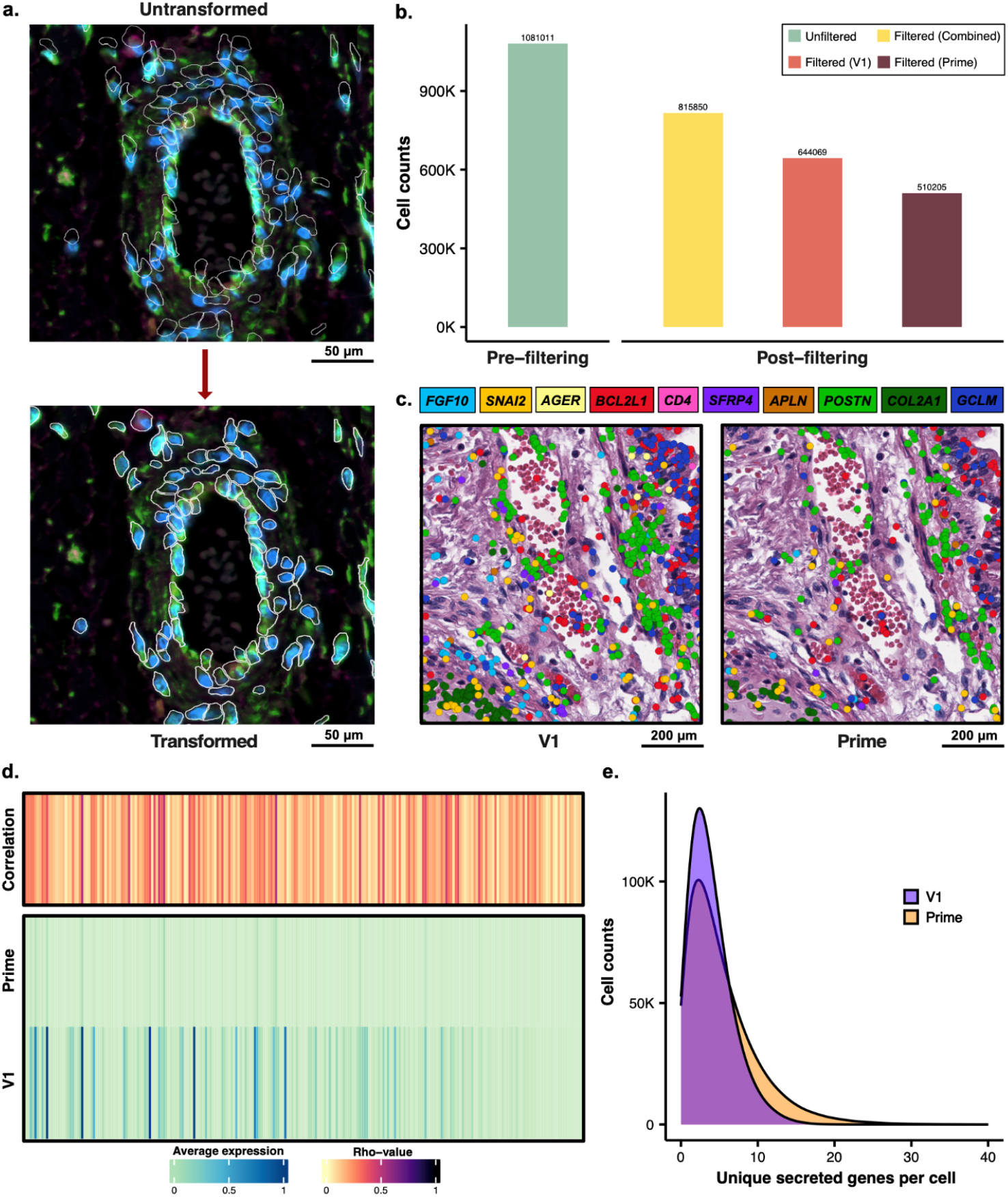
Spatial Linear Transformation, Quality Control Filtering, and Visualization of Transcript Distribution in Dual Runs. a) The top panel displays the untransformed dual Prime run image with the original V1 cell segmentation masks (white outlines). The bottom panel displays the result after applying a linear transformation to correct for subtle shifts, successfully aligning the segmentation masks with the underlying cellular morphology. b) Bar chart comparing the initial total cell count (pre-filtering) of the dual run to the final cell counts (post-filtering) achieved after applying three layers of filtering to three distinct gene sets: the combined gene panel (yellow), the V1-only gene panel (orange), and the Prime-only gene panel (brown). c) Two adjacent panels show the same histological field of view, with underlying morphology by H&E staining. The left panel overlays the spatial distribution of transcripts detected by the V1 panel, and the right panel overlays the transcripts detected by the Prime panel. Both images display the expression of 10 selected genes, with their respective transcripts colored according to the legend above. d) A heatmap quantifying the relationship between the V1 and Prime panels by showing the average expression of 239 overlapping genes and their expression correlation (Rho-value). e) A density plot that compares the efficiency of the panels in capturing unique secreted genes per cell, demonstrating that the Prime panel (538 genes) identifies a higher number of unique secreted genes than the V1 panel (59 genes).

Given the shared 239 genes in the V1 and Prime panels, we were able to perform cross chemistry comparisons. First performing a qualitative assessment of 10 representative overlapping genes we observed similar expression profiles for the V1 and Prime Chemistries across differing architectures – albeit with lower transcript counts in the Prime chemistry (Fig. 2c, Fig. S5). Nevertheless, we observe only modest correlations between the V1 and Prime expression levels on the dual (see Methods; Fig. 2d; Table S7). Indeed, the low correlation is likely driven by the differences in sensitivity between the two chemistries (Fig. 2d). A large portion of cells in the dual run dataset have a low dynamic detection range in Prime, with many observed zeros. Furthermore, when we compared the overall slide level correlation for the 239 genes on both the V1 and Prime chemistries, we see very similar trends in both the dual and solo runs (Fig. S6). Together this suggests the modest correlations between the dual V1 and Prime data are driven by technical differences inherent to the chemistry – rather than the dual run approach. However, the true value of this integrated approach is the ability to couple a targeted gene panel – focused on cell type annotation and known biological markers – with a more discovery-oriented approach. While this study was not designed to showcase the breadth of these possibilities, an example of such an analysis is illustrated by the “secretome”. Focusing on the expression of genes that are known to be secreted, we first observe an obvious imbalance in the panels with 538 secreted genes profiled with the Prime chemistry and only 59 profiled on V1. Furthermore, when we compare the number of unique secreted genes observed in each cell (requiring at least 1 count) we observed a higher number in the prime chemistry – with some cells having over 20 unique secreted genes (Fig. 2e; Tabs. S2, S3, S8). This illustrated the increased discovery potential for diverse secretory activity and unique secretory signals within our samples. At this number of genes, the same analysis would have required that our entire 480 gene V1 panel be dedicated only to secretory markers in order to profile the same diversity of markers.

Finally, the additional information provided by having both panels exist in the same cells enhances the potential of downstream analyses. In this study the V1 panel was custom designed for cell typing accuracy and biological relevancy in the human lung. Thus, we observe better resolution, more clusters, and more retained cells when looking at the V1 only compared to the Prime. However, when combining these datasets for a unified analysis (see Methods), we were able to recover more cells, while retaining similar cluster resolution (Fig. 3a). While we observed somewhat stable cluster clusters (Fig. 3b), there was higher concordance between the dual and V1 than the V1 and Prime. However, given that the V1, prime, and dual datasets are all generated from the same cells, researchers can choose the clustering approach that is best suited for their projects (Fig. S7). Together, these results support the idea that the dual Xenium Prime and V1 prep allows for an increased depth and breadth of transcript information available at the cellular level.

**Figure 3:**
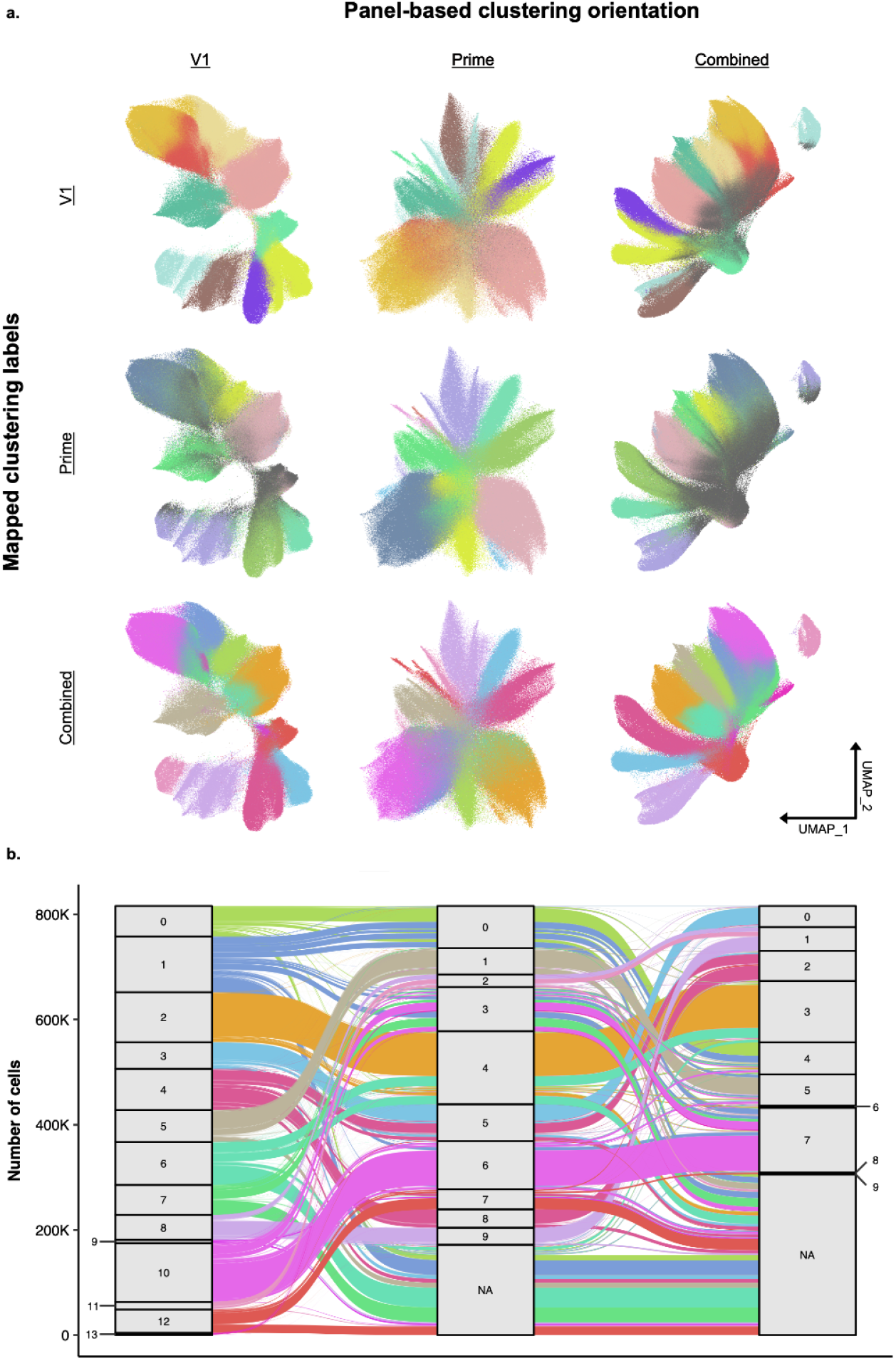
Tracing Cell Fate: Cluster Transition and Stability Between V1, Prime, and Combined Panels. a) A matrix of UMAPs comparing three panel-based clustering orientations (V1, Prime, and Combined across the x-axis) against three sets of mapped clustering labels (V1, Prime, and Combined across the y-axis). This illustrates the degree of cluster structure preservation when mapping labels from one clustering method onto another’s UMAP space. The cells colored in gray were filtered out per that panel (e.g., the V1 UMAP with Combined cluster labels has a small portion of grayed-out cells, indicating that those cells were filtered out with V1-panel specific filtering metrics but retained with combined-panel specific metrics). b) An alluvial plot that highlights the distribution and transitions of cell labels across clustering resolutions (Combined, V1, and Prime). The plot shows the number of cells belonging to each cluster (stratum) and how those populations flow between the different clustering results, highlighting the stability and splitting of clusters.

## Discussion

There are several limitations to the work described in this study. First, the sample set contained in this paper is limited to a single group of 17 unique pulmonary samples. We are not able to draw any conclusions about the replicability of the results observed across other tissue types. We were able to use the two adjacent sections prepared with the V1 and Prime solo chemistries to judge performance, but even those are imperfect proxies. Secondly, the fact that V1 regularly outperforms Prime for sensitivity can make the comparison a bit difficult to judge performance between the solo chemistries and the dual. We attempted to mitigate this by looking largely at the dual slide as a single dataset rather than assessing each chemistry separately. This is also largely the reason that the V1 chemistry results in better filtering and clustering, so the relative comparisons to dual are not as easily discerned.

A current barrier for widespread implementation of this method is the increased cost per slide. In order to prepare the data in this paper, the dual prep required the purchase of both a V1 Xenium Sample Prep Reagents kit (10X Genomics 1000460) and a Xenium Prime 5K Human sample prep reagents kit (10X Genomics 1000671). After the completion of this data collection, and through discussion with the R&D team at 10X Genomics, we were made aware of a potential protocol alteration that could lessen this burden. This adaptation may enable dual detection of a V1 panel and the Prime panel with the use of only the Prime sample prep reagents. It does require the use of an additional oligo of sequence **GGCTCCACTAAATAGACGCA** and resuspending it at 100uM. The resuspension should follow the steps used for custom Xenium panels, but adjust the volume to obtain the desired 100uM based on the purchased oligo yield. The panels should be mixed as described in the methods here and hybridized accordingly. While performing the dual prep Xenium ligation steps, instead of using the Xenium Ligation Buffer (10X Genomics, 2000391) from the V1 chemistry kit, use the Ligation Buffer B (10X Genomics, 2001233) from the Prime chemistry kit and spike in 1uL of the resuspended oligo into 1000uL of Ligation Buffer B for a final oligo concentration of 100nM. This Ligation Buffer B can then be mixed as normal with the ligation enzymes and continue the prep as described. As this protocol adaptation has not been tested in our hands, we cannot speak to its performance, but wanted to make the reader aware of its potential. If the results are similar to those seen here, it could be a viable option to slightly reduce the monetary cost of this dual detection method.

Finally, while we only provide one example of the opportunities provided by having a combined dataset, we believe there are significant opportunities for discovery. These range from obvious additional analyses, such as ligand receptor analysis carried out on the background of high-fidelity cell typing, to more nuanced approaches, such as deriving project specific imputation approaches – using the V1 data to impute the missing Prime or vice versa. Indeed, an imputation approach would enable researchers to only perform the dual profiling on a subset of samples. Finally, while the sensitivity benefits of small custom probe-based studies have been appealing, they suffer from difficulties with cross-study integration. Since the advent of genomics, the field has benefited from mappings to common references that allowed large post-hoc meta-analyses. However, with small curated panels of genes, even studies within the same organ system may have limited overlap – depending on the motivation of the research. This severely hinders the ability to integrate disparate datasets leading to a diminished future utility. In contrast, this approach provides a stable common reference (the Prime chemistry) without sacrificing sensitivity of key genes. This shared target space will allow cross-study integration providing future value for the scientific community.

The ability to co-detect tissue and sample-specific V1 panels alongside Prime 5K panels in the same cells without signal dropout opens the door to greater biological understanding. It provides detection in a large shared 5001 gene base on which large datasets across institutions can be aligned, while also allowing for individualized V1 panels to target specific questions or discoveries. Having greater combined histological and expression context is the central goal of Xenium spatial transcriptomic technology, meaning that having the breadth of the Prime 5K panel matched with the depth and sensitivity of a V1 panel maximizes that potential as far as the current technology allows.

## Methods

### Sample selection

Samples included in this study were archived formalin fixed paraffin embedded (FFPE) tissue from human lung. A section was taken from the face of each sample block for standard H&E staining. A team with expertise in pediatric lung pathology selected a 3mm × 3mm target location for the region of interest.

### Tissue microarray (TMA) construction

The Xenium capture area is 10.45mm × 22.45mm. The use of a tissue microarray (TMA) allows for the more precise placement of multiple tissue samples within that area. The FFPE TMA block was designed in a 3 × 6 pattern of 17 3mm cores, with the top left core space remaining empty as an orientation marker.

A custom 3 mm square tip was created by the Arizona State University Instrument Design and Fabrication Core to fit the commercially-available Quick-Ray Manual Tissue Microarrayer and allow us to produce square cores. The TMA was constructed by placing each core individually, tissue face down, on double-sided adhesive tape in a paraffin block mold^7,8^. The entire mold is warmed to 37 °C, and a P1000 is used to dispense melted paraffin wax in the junctions between cores. More wax is poured over the back of the block to fill in the remaining space and adhered in a plastic cassette. Once solidified, the block was extracted from the mold, tape was removed, and the block was stored at 4°C until sectioning.

### Sectioning and sample placement

The Leica RM2135 microtome and surrounding area was cleaned with RNase AWAY and 70% Isopropanol prior to sectioning. The TMA block was rehydrated using nuclease-free water and then cut with a DURAEDGE Low-Profile PTFE blade at 5um. Sections were floated on a 40°C heated water bath filled with nuclease-free water, to help remove wrinkles. Three serial sections were taken and each placed on an individual Xenium slide. Slides dried overnight at room temperature, then baked the next day for 3 hours at 37°C. Following this incubation, slides were stored in a sealed desiccator at room temperature for 14 days prior to Xenium sample preparation.

### Gene panel design

All Xenium runs, V1 or Prime 5K, require the use of a gene panel. A total of 480 genes were included in our V1 custom panel. These genes were chosen to cover expression in the human lung and enable cell-type identification based on single-cell analysis data^9^ and Xenium analysis data^10^. The Xenium Prime 5K Human Pan Tissue and Pathways panel contains 5001 genes and was designed by 10X Genomics to cover expression in a variety of human tissues. 239 genes are present on both our custom V1 panel and the predesigned Prime panel (Tabs. S2 - S3).

### Custom probe resuspension

New tubes of our custom V1 panels were used for this experiment. This facilitated the resuspension of the lyophilized probes in a lower volume than normal, which is required to properly spike into the Prime panel and maintain relative concentration. This also allowed both the V1 only slide and the dual slide to be prepped with the same lot of probes to maintain consistency.

6ZAAWG, our custom base panel, was ordered in a 16 reaction quantity and resuspended in ⅓ of the normal volume TE Buffer (233.33uL versus 700uL, 10X Genomics CG000749). 6ZX7JP and 43R3RN, our custom add-on panels, were ordered in 16 reactions quantities and each resuspended in ⅙ of the normal volume TE buffer (116.67uL versus 700uL, 10X Genomics CG000749). In a normal V1 run, these 50 gene panels are still resuspended at double the concentration of a single add-on in order to add both to 6ZAAWG while maintaining the 33uL add-on volume (16.5uL each).

### Xenium sample preparation

Xenium relies on intact in situ RNA for binding, so all workstations and equipment are cleaned using RNase AWAY (VWR, 53225-514) and followed by 70% isopropanol. All reagents, including water, were molecular-grade nuclease-free to further preserve RNA quality.

The V1 only slide underwent deparaffinization and decrosslinking according to the 10X Genomics Demonstrated Protocol CG000578 without alteration. This slide underwent probe hybridization using the following probe mix recipe: 315uL Xenium Probe Hybridization Buffer, 144uL TE Buffer, 33uL 6ZAAWG, 16.5uL 2X 6ZX7JP, 16.5uL 2X 43R3RN. Probe hybridization occurred at 50°C for 19 hours. Ligation, amplification, cell segmentation, autofluorescence quenching, and nuclei staining were performed according to the 10X Genomics User Guide CG000749 without alteration. Slide was stored at -20°C in 30% glycerol solution in PBS for 18 days following sample prep completion before loading onto the Xenium Analyzer.

The Prime only slide underwent deparaffinization and decrosslinking according to the 10X Genomics Demonstrated Protocol CG000578 without alteration. This slide underwent probe hybridization using the following probe mix recipe: 94.5uL Probe Hyb Buffer B, 10.5uL TE Buffer, 52.5uL Xenium 5K Hu PTP Panel Probes. Probe hybridization occurred at 50°C for 16 hours. Priming, RNase treatment, polishing, ligation, amplification, cell segmentation, autofluorescence quenching, and nuclei staining were performed according to the 10X Genomics User Guide CG000760 without alteration. Slide was stored at -20°C in 30% glycerol solution in PBS for 22 days following sample prep completion before loading onto the Xenium Analyzer.

The Dual chemistry slide underwent deparaffinization and decrosslinking according to the 10X Genomics Demonstrated Protocol CG000578 according to the Prime chemistry directions. This slide underwent sample preparation according to the 10X Genomics User Guide CG000760 Prime chemistry with the following alterations. The probe hybridization mix recipe: 94.5uL Probe Hyb Buffer B, 52.5uL Xenium 5K Hu PTP Panel Probes, 5.25uL 3X 6ZAAWG, 2.63uL 6X 6ZX7JP, 2.63uL 6X 43R3RN. Probe hybridization occurred at 50°C for 16 hours. For the ligation steps use Xenium Ligation Buffer (10X Genomics, 2000391) from the V1 chemistry kit instead of Ligation Buffer B (10X Genomics, 2001233) from the Prime chemistry kit. At the end of the prep, the slide was stored at -20°C in 30% glycerol solution in PBS for 18 days before loading onto the Xenium Analyzer for the first run.

### Xenium Analyzer imaging

Detailed instructions for instrument configuration, consumables, and buffer preparation can be found in 10X Genomics User Guide CG000584. After loading and manual area selection on a low-resolution full-slide image, the Xenium Analyzer fully automates the collection of fluorescent puncta data through rounds of fluorescent probe binding, imaging, and stripping. Each potential RNA puncta location is then decoded and labeled by gene ID, according to the fluorescence pattern across imaging channels and cycles. All image fields of view (4,240 × 2,960 pixels) are then stitched together using the nuclei-stained DAPI image with RNA transcript locations overlaid in the x-y-z coordinate system of the images. Onboard instrument analysis provides quality metrics for each detected transcript based on decoding confidence from the image cycle signals. Additional fluorescent images are taken at the end of the run to enable cell segmentation based on cell nuclei, cell boundary, RNA, and interior protein staining. All data in this paper were collected on instrument software version 3.4.1.0 and onboard analysis version xenium-3.3.0.1.

The Xenium Analyzer instrument performs data acquisition for a maximum of two slides per run. First loaded were the V1 only and the dual slide for a V1 decoding run. Cell segmentation was performed on this run. Following run completion, slides were removed from the instrument. The post-run buffer in the slide cassette wells was removed and replaced with fresh 1000uL PBS-T. The V1 slide was stored at 4°C until post-run histology staining was performed. The dual slide was stored at 4°C for ∼12-16 hours before reloading onto the instrument for a second run.

The second batch of slides loaded were the prime only and the dual slide for a prime 5K decoding run. Cell segmentation was performed on this run. Following run completion, slides were removed from the instrument. The post-run buffer in the slide cassette wells was removed and replaced with fresh 1000uL PBS-T. Both slides were stored at 4°C until post-run histology staining was performed.

### Post-Xenium histology

All three slides underwent quencher removal in sodium hydrosulfite according to 10X Genomics Demonstrated Protocol CG000613. Immediately following, the slides were H&E (hematoxylin and eosin) stained according to the following protocol: DI water (2 min), Mayer’s hematoxylin (2 min; Millipore Sigma MHS16-500ML), DI water (x3, 1 min each), bluing solution (1 min; Agilent CS70230-2), DI water (1 min), 95% ethanol (3 min), eosin Y (45 s; Fisher Scientific NC2101164), 95% ethanol (1 min), 100% ethanol (x2, 30 s each), xylene (x2, 3 min each). Coverslipping was performed using ∼150uL Leica Mounting Media (Fisher Scientific NC1109222) and #1.5 50x24mm cover glass (VWR 16002-264). Mounting media was cured for at least 30 minutes at room temperature. Histology images were taken on a 40X (20X objective with doubler inserted) Leica Biosystems Aperio CS2.

### Cell segmentation

Cell segmentation was performed on the Xenium Analyzer instrument using the onboard multimodal cell segmentation algorithm with default parameters (onboard analysis version xenium-3.3.0.1).

### Image registration

Image registration between the DAPI images of the V1 and Prime dual runs was performed in the Xenium Explorer software (version 4.1.0). Briefly, the DAPI image from the V1 run was aligned to that of the Prime run through a linear transformation based on the placement of anchors on corresponding landmarks in both images.

### Cell mask transformation

Cell masks from the V1 run were transformed into the coordinate system of the Prime run using the linear transform produced by image registration and the import-segmentation command within Xenium Ranger (version 4.0). Final fine-scale adjustments to the alignment were made using a custom Python script. After these adjustments were made, import-segmentation was run again to update the transcript assignments with this final alignment.

### Quality control - upstream filtering

Raw Xenium output resulted in transcript and cell metadata information. Each transcript file (per slide) contained metrics for each transcript, including spatial coordinates, cell-level information (cell_id), quality value (QV), and nuclear overlap (overlaps_nucleus). We used Seurat^11^ to create objects with this information and filtered transcripts to remove any that did not meet quality parameters: QV < 20, overlaps_nucleus = 0, and cell_id = UNASSIGNED. We created gene matrices based on the remaining transcripts that were partitioned into a segmented nucleus, and with those matrices and metadata information, generated Seurat objects per each slide that were merged into a singular object.

### Panel-specific object creation

One Seurat object was made with the output information from the V1 solo run and the dual run with V1 segmentation, and one with the prime solo run and the dual run with prime segmentation. The dual V1/Prime run contains the same cells across both iterations. However, Xenium utilizes randomly-generated strings for cell labels, and to ensure that the cells are correctly matched, the unique string was generated per row of metadata across the run with the V1 segmentation and the prime segmentation to match cells accordingly post-linear transformation. In order to successfully combine the gene matrices across the V1 and prime panels, unique identifiers were appended to each feature name in the transcript files, which would allow the matrices to be combined into one.

We then used Scanpy^12,13^ to create an Anndata object with only the dual run, but both segmentation types, from these adjusted transcript data. We calculated various metrics (number of counts/genes per cell) with respect to the V1, prime, and the combined panel genes. We then created two following Anndata objects - one with the V1 and prime genes respectively. Cells from each object were retained according to the following parameters: n_counts >= 50, n_genes >= 5, cell_area >= 5 & <= 140, nucleus_area >= 3.

### Clustering

To do any clustering, we required the use of a graphical processing unit, given the magnitude of data. We used a container with rapids_singlcell^14^ - single cell Python library (ScanPy - v1.8.1) to perform clustering and dimensionality reduction across all three panel-specific objects. Each object was clustered with 20 PCs and 15 nearest neighbors.

### Correlations

In this study, we compared gene expression 239 overlapping genes across the V1 and prime panels using a Spearman’s correlation test. To do this, we aggregated the expression across all cells per gene into one value. From there, the correlation was done between each shared gene across both panels.

## Supporting information

Main Tables and Figures

## Protocol availability

Step-wise methods can be found on protocols.io at DOI: dx.doi.org/10.17504/protocols.io.5qpvodwm7g4o/v1.

## Data availability

Raw and processed data are deposited at GEO under accession **GSE315411**.

## Code availability

Custom R and Python scripts for this project are available on GitHub at https://github.com/Banovich-Lab/xenium_v1_prime_comparison.

## Funding

This work was supported by National Institutes of Health (NIH) U01HL175444 (to J.A.K., N.E.B., and J.M.S.S.), R01HL145372 (to J.A.K and N.E.B), and R01HG011886 (to N.E.B). Lung samples were collected under a study approved by the Institutional Review Board for human subject research at Seattle Children’s Hospital, STUDY00003664 (G.H.D.).

## Notes

http://dx.doi.org/10.17504/protocols.io.5qpvodwm7g4o/v1

https://github.com/Banovich-Lab/xenium_v1_prime_comparison

### Competing Interest Statement

The authors have declared no competing interest.

