## Supplemental Figures for "Fishing with Two Lines: A Hybrid Approach to Spatial Transcriptomic Discovery"

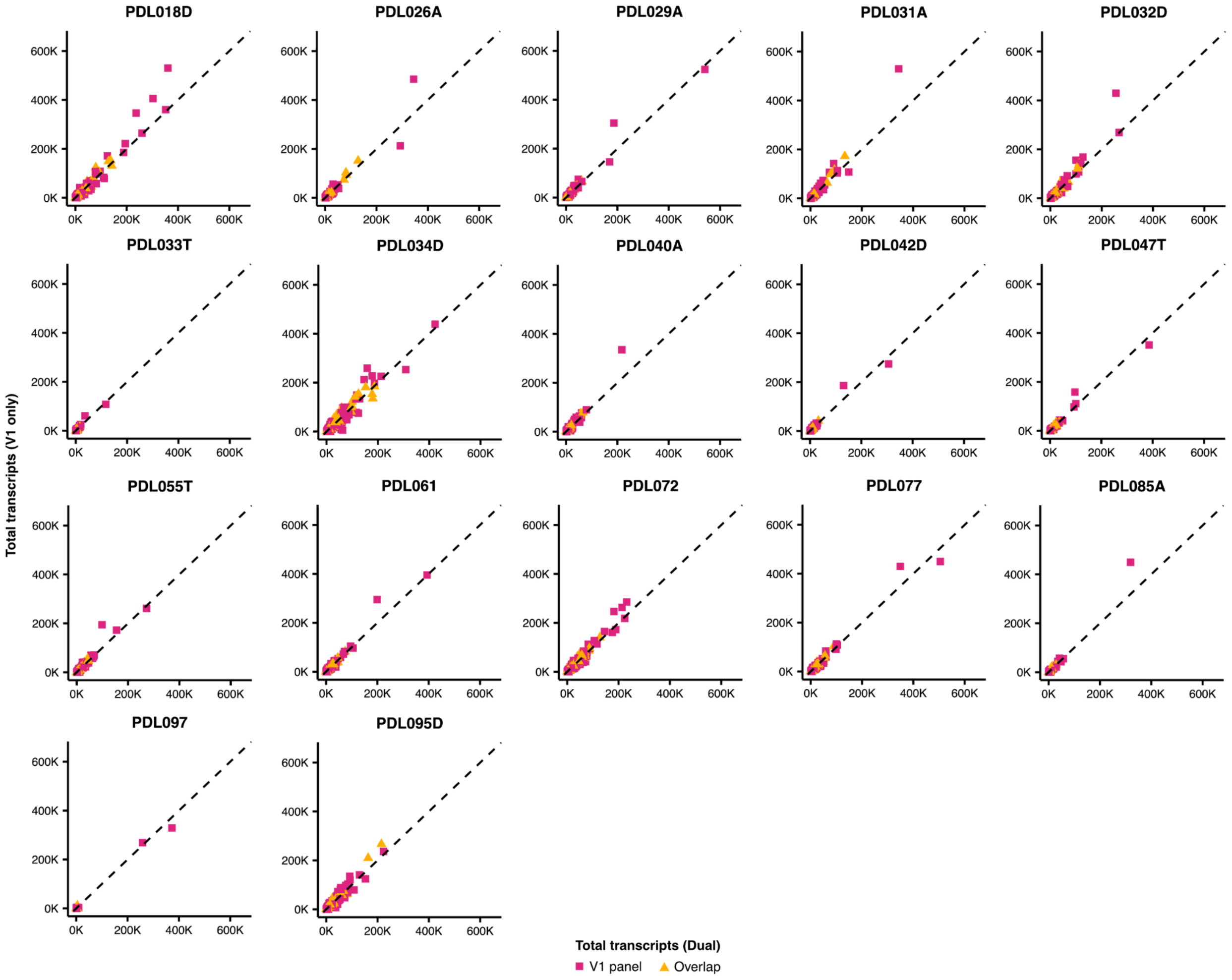


**Figure S1:** Scatterplots that show the correlation of raw per-gene transcript counts across all genes detected in the V1 panel, split by sample. The x-axis represents the total transcripts for the V1 panel genes, and the y-axis represents the total transcripts for the dual run genes.

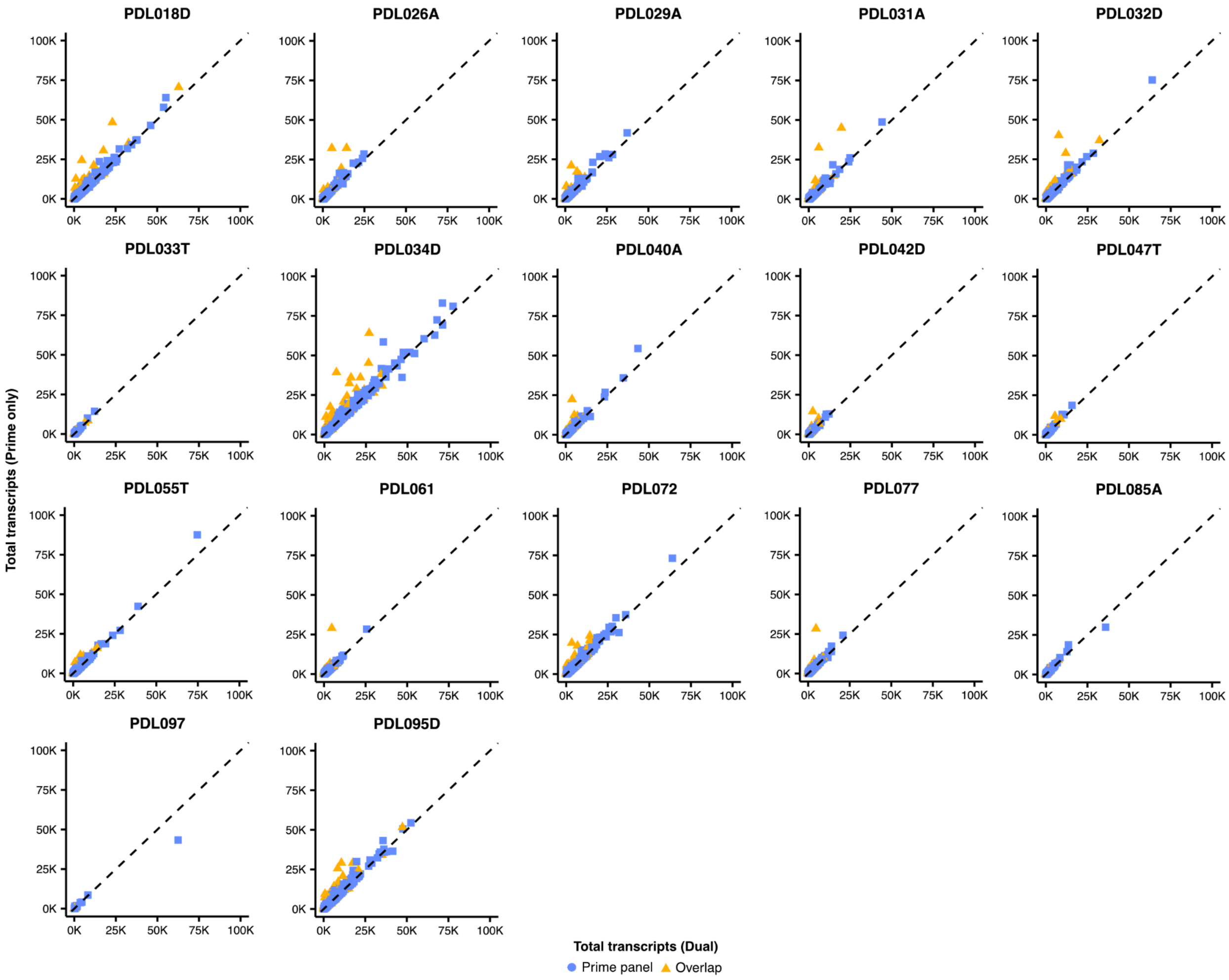


**Figure S2:** Scatterplots that show the correlation of raw per-gene transcript counts across all genes detected in the Prime panel, split by sample. The x-axis represents the total transcripts for the V1 panel genes, and the y-axis represents the total transcripts for the dual run genes.

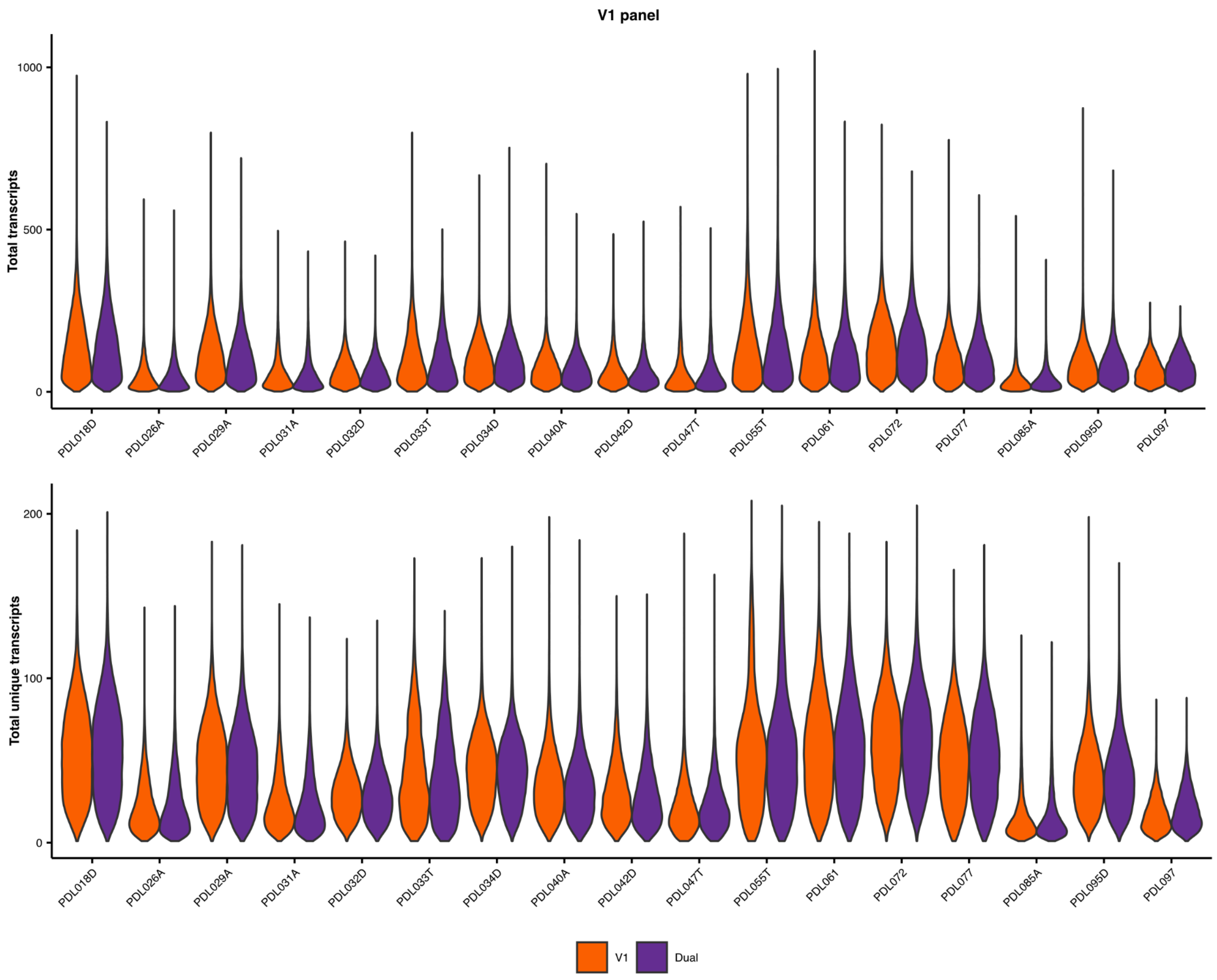


**Figure S3:** Violin plots that display the distribution of total transcripts and total unique transcripts for V1 and the combined dual run, split by sample.

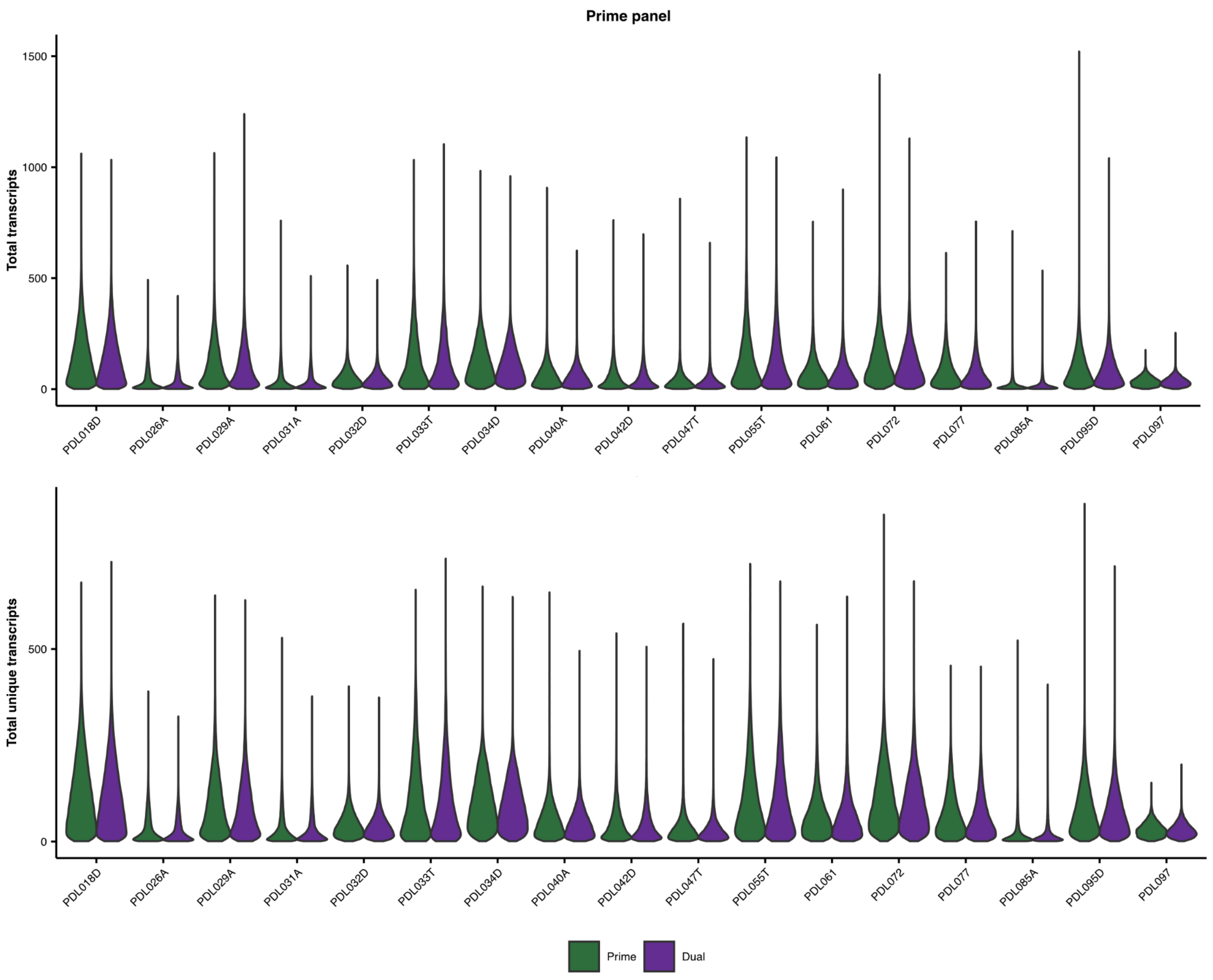


**Figure S4:** Violin plots that display the distribution of total transcripts and total unique transcripts for Prime and the combined dual run, split by sample.

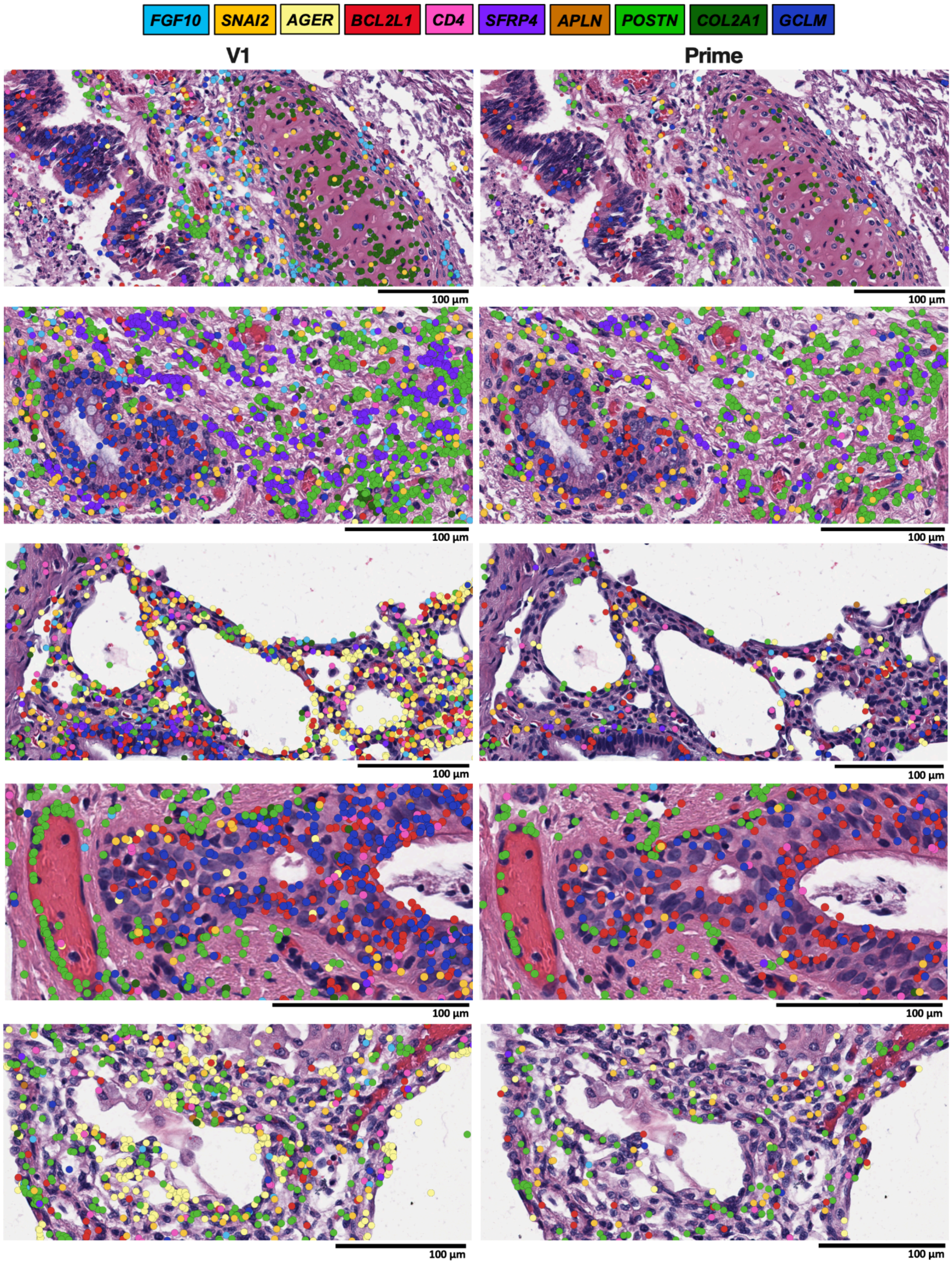


**Figure S5:** Additional histological fields of view showcasing the spatial distribution of V1 gene detection (left) and Prime gene detection (right), with underlying H&E morphology. Both images display the expression of 10 selected genes, with their respective transcripts colored according to the legend above.

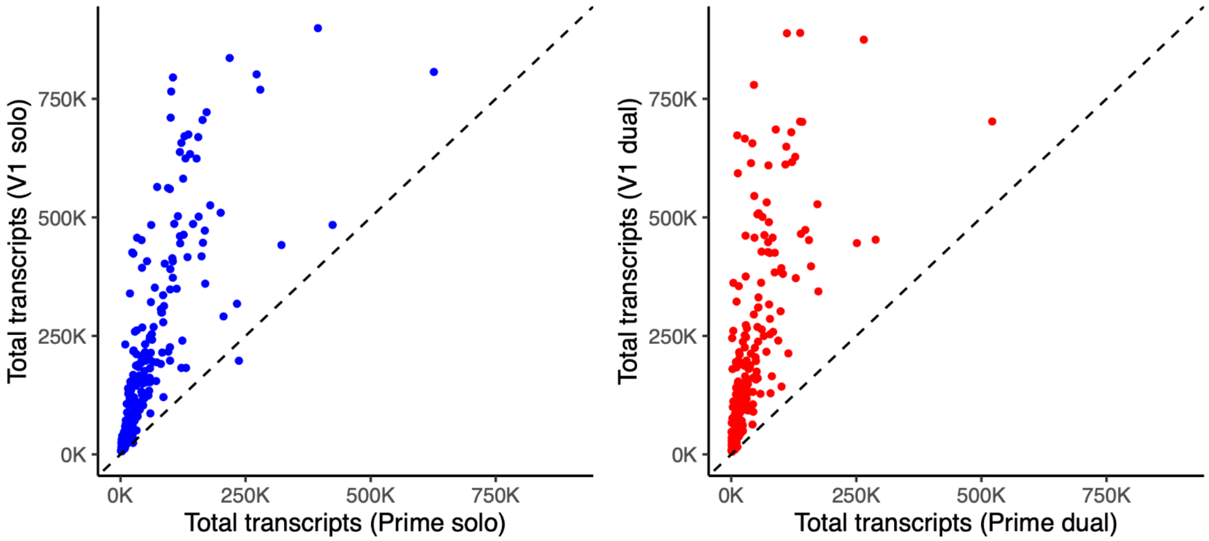


**Figure S6:** Scatterplots that show the correlation between total detected transcript counts between V1 and Prime chemistries run alone and then both chemistries on the dual slide. Each point represents the transcript count for one of the 239 overlapping genes.

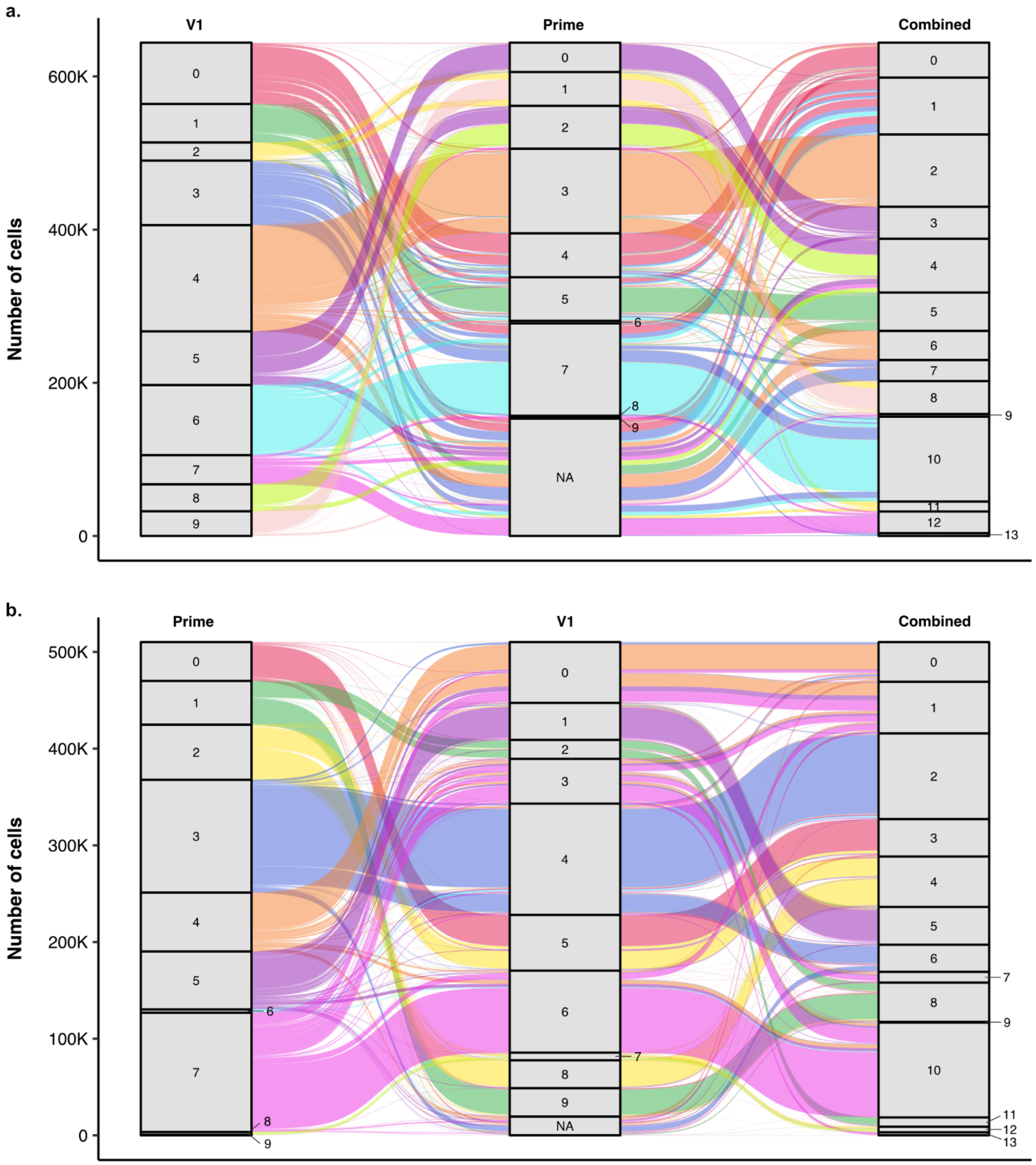


**Figure S7:** a) An alluvial plot that highlights the distribution and transitions of cell labels across clustering resolutions (V1 to Prime to Combined). b) The same plot but in the order of Prime to V1 to Combined.
