## Supplementary material for "Fishing with Two Lines: A Hybrid Approach to Spatial Transcriptomic Discovery": Main Tables and Figures

| | Solo V1 | Dual V1 | Dual 5K | Solo 5K |
|---|---|---|---|---|
| **Number of Cells Detected** | 1,111,107 | 1,107,543 | 1,115,068 | 1,100,358 |
| **Median Transcripts per Cell** | 125 | 123 | 100 | 115 |
| **Nuclear Transcripts per 100um$^2$** | 330.8 | 322.3 | 278.8 | 314.5 |
| **Total Transcripts** | 221,439,804 | 218,969,052 | 187,967,974 | 210,218,833 |

**Table 1: Slide-level transcript detection metrics.** A table containing per slide segmented cell numbers, median transcript count per cell, average transcripts overlapping segmented nuclei per 100um$^2$, and the total transcripts detected across the full slide.

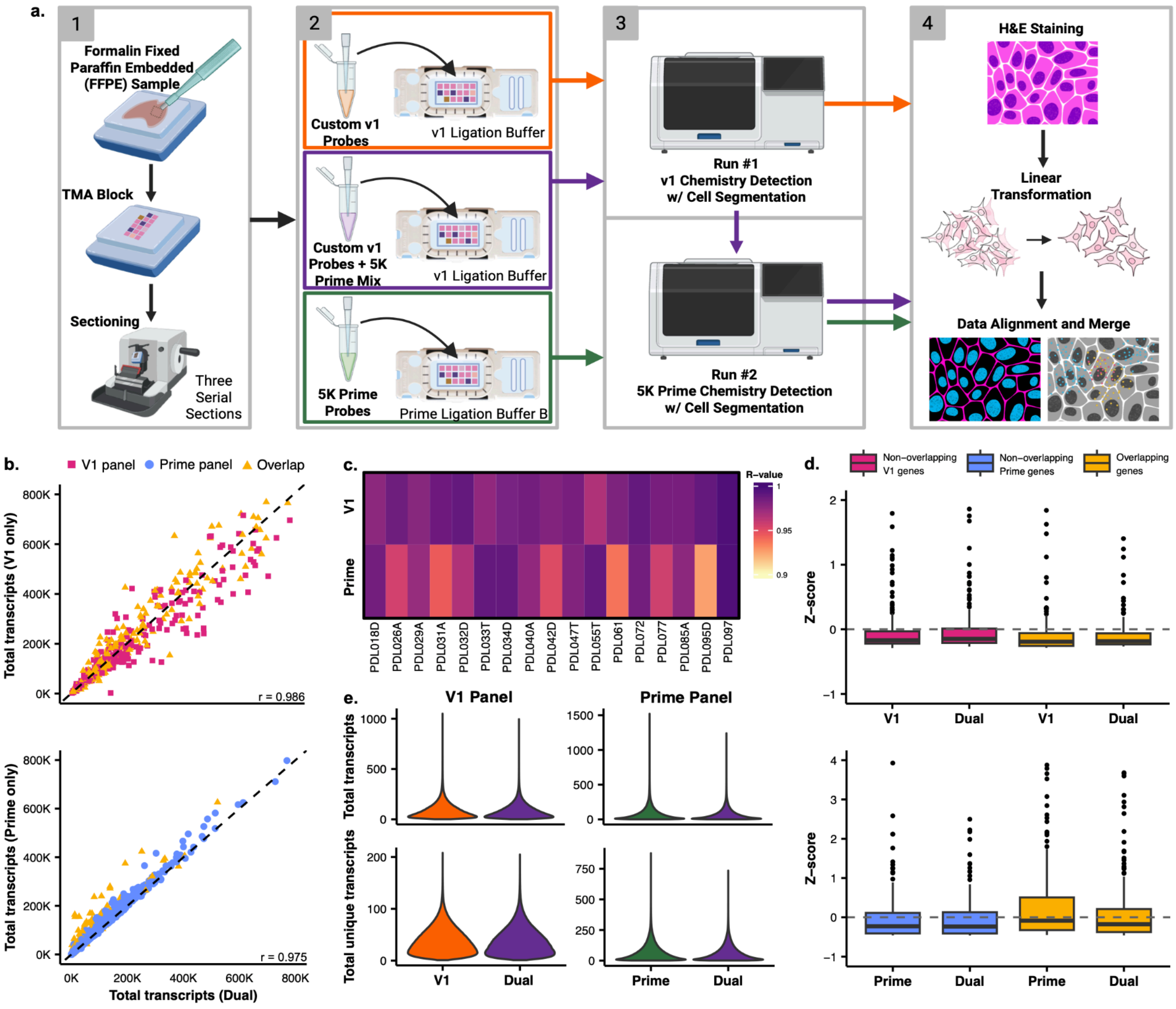


**Figure 1: Evaluation of Transcript Quantification Consistency and Performance Across V1 and Prime Panel Assays.** a) Schematic representation of the study design. b) Scatterplots that show the correlation of raw per-gene transcript counts across all genes detected in the V1 panel (top) and the Prime panel (bottom). The x-axis represents total gene expression from the dual assay (V1 + Prime or Prime + V1), while the y-axis represents total gene expression from the solo assay (V1 only or Prime only). Genes are colored by panel specificity: V1-specific genes (pink), prime-specific genes (blue), and genes overlapping both panels (yellow). c) Heatmap visualizing the R-value derived from correlating the transcript counts from the solo runs with the dual-run transcript counts for 17 infant lung samples. d) Boxplots that display the z-score distribution for three gene sets: the 239 genes overlapping between the V1 and Prime panels, the non-overlapping V1-specific genes, and a random subset of 239 non-overlapping Prime-specific genes. Genes are colored by their group: Overlapping (yellow), non-overlapping V1-specific (pink), and non-overlapping prime-specific (blue). e) Violin plots of the distribution of total transcripts and total unique transcripts for the three main run types: V1 only, prime only, and the combined dual run.

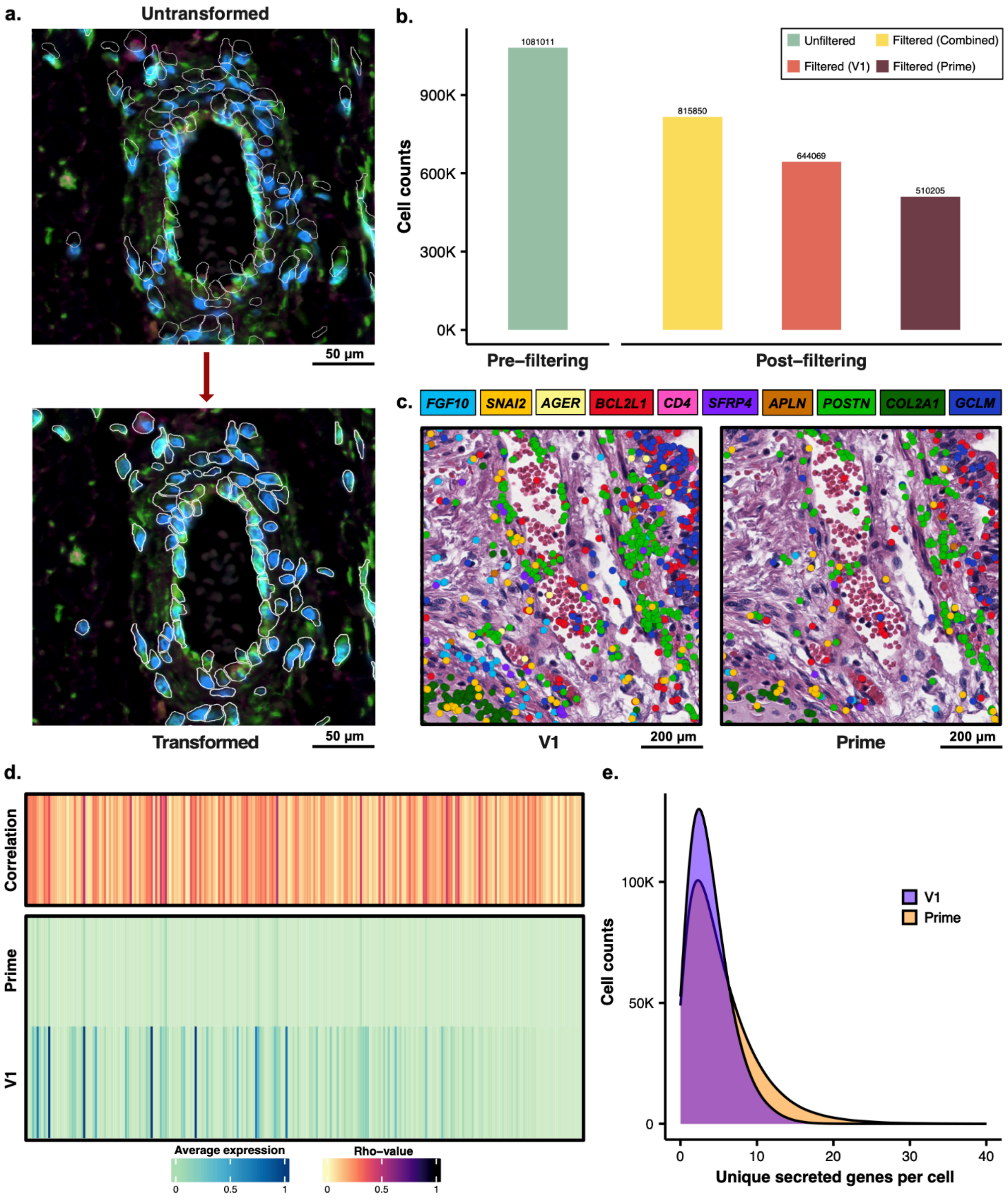


**Figure 2: Spatial Linear Transformation, Quality Control Filtering, and Visualization of Transcript Distribution in Dual Runs.** a) The top panel displays the untransformed dual Prime run image with the original V1 cell segmentation masks (white outlines). The bottom panel displays the result after applying a linear transformation to correct for subtle shifts, successfully aligning the segmentation masks with the underlying cellular morphology. b) Bar chart comparing the initial total cell count (pre-filtering) of the dual run to the final cell counts (post-filtering) achieved after applying three layers of

filtering to three distinct gene sets: the combined gene panel (yellow), the V1-only gene panel (orange), and the Prime-only gene panel (brown). c) Two adjacent panels show the same histological field of view, with underlying morphology by H&E staining. The left panel overlays the spatial distribution of transcripts detected by the V1 panel, and the right panel overlays the transcripts detected by the Prime panel. Both images display the expression of 10 selected genes, with their respective transcripts colored according to the legend above. d) A heatmap quantifying the relationship between the V1 and Prime panels by showing the average expression of 239 overlapping genes and their expression correlation (Rho-value). e) A density plot that compares the efficiency of the panels in capturing unique secreted genes per cell, demonstrating that the Prime panel (538 genes) identifies a higher number of unique secreted genes than the V1 panel (59 genes).

a.

**Panel-based clustering orientation**

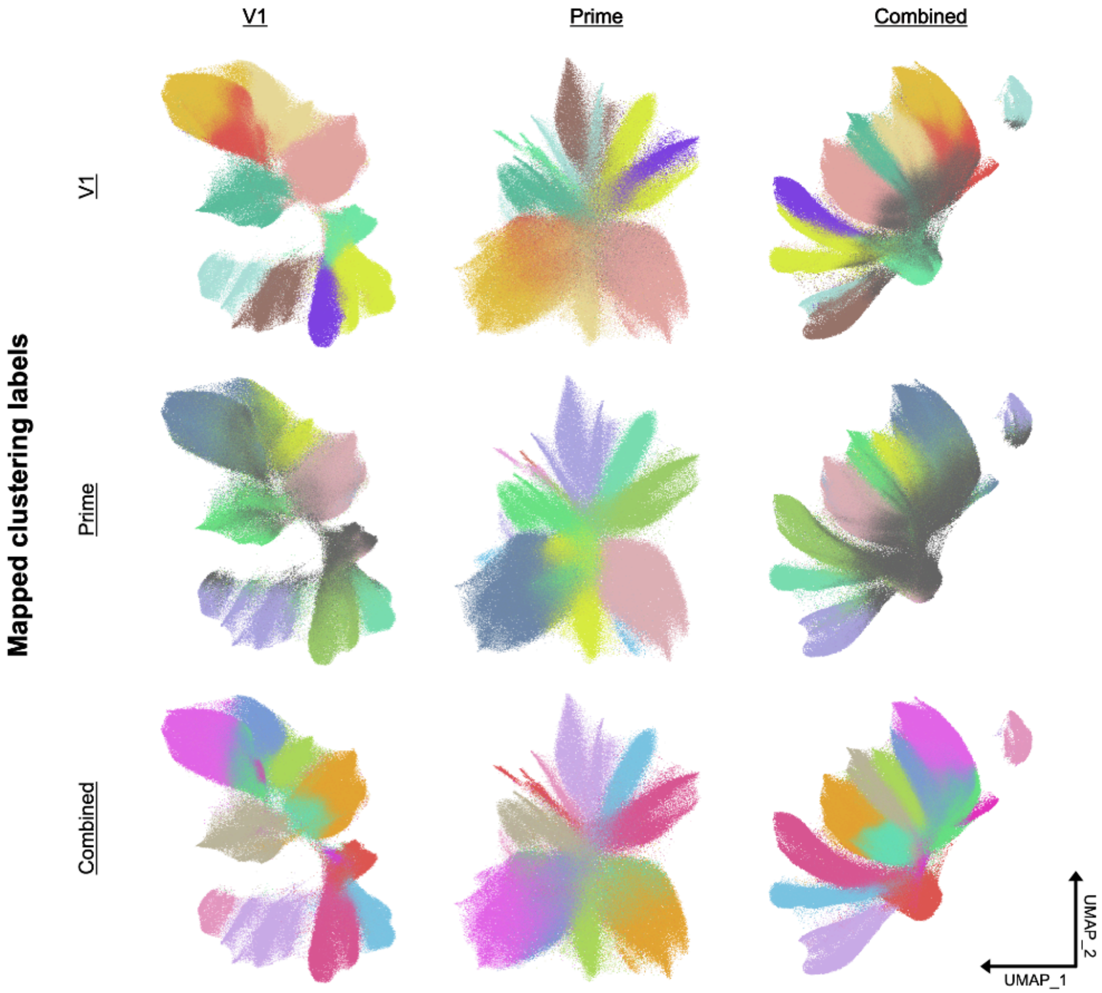


b.

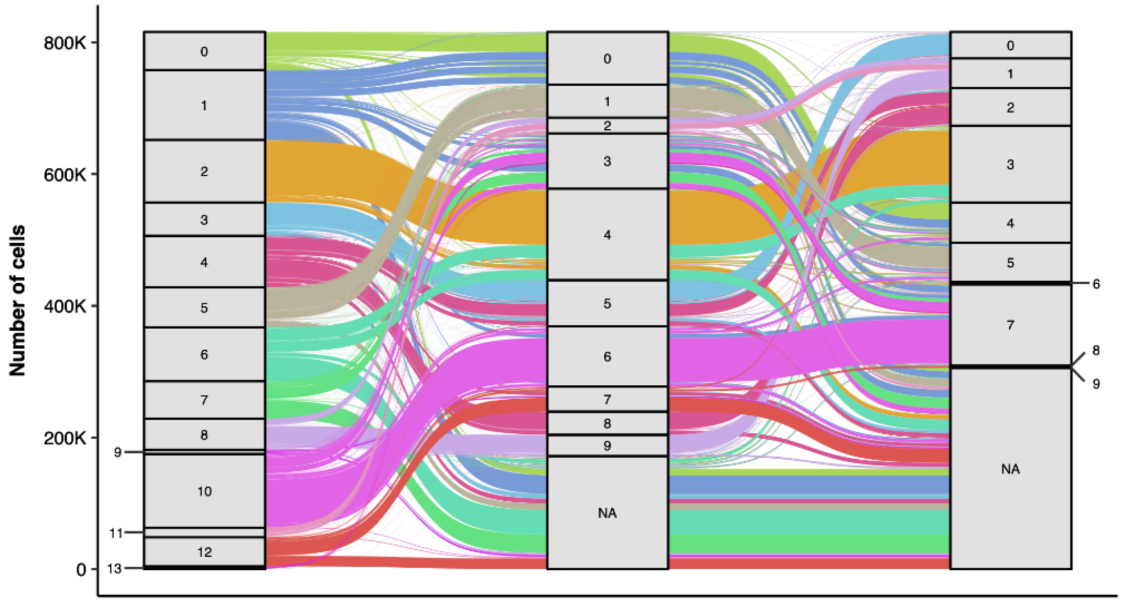

**Figure 3: Tracing Cell Fate: Cluster Transition and Stability Between V1, Prime, and Combined Panels.** a) A matrix of UMAPs comparing three panel-based clustering orientations (V1, Prime, and Combined across the x-axis) against three sets of mapped clustering labels (V1, Prime, and Combined across the y-axis). This illustrates the degree of cluster structure preservation when mapping labels from one clustering method onto another's UMAP space. The cells colored in gray were filtered out per that panel (e.g., the V1 UMAP with Combined cluster labels has a small portion of grayed-out cells, indicating that those cells were filtered out with V1-panel specific filtering metrics but retained with combined-panel specific metrics). b) An alluvial plot that highlights the distribution and transitions of cell labels across clustering resolutions (Combined, V1, and Prime). The plot shows the number of cells belonging to each cluster (stratum) and how those populations flow between the different clustering results, highlighting the stability and splitting of clusters.
